## Supplementary Figures S1-S6 for "A CENH3 mutation promotes meiotic exit and restores fertility in SMG7-deficient Arabidopsis"

**Supplementary Movies and Figures**

**Movie S1: Live imaging of *pRPS5A::TagRFP:TUB4* tubulin marker during the first and second meiotic division in wild a type plant.**

**Movie S2: Live imaging of *pRPS5A::TagRFP:TUB4* tubulin marker during regular meiotic divisions and two postmeiotic spindle reassembly cycles in smg7-6 mutants.**

**Movie S3: Live imaging of *pRPS5A::TagRFP:TUB4* tubulin marker from the 3$^{rd}$ to 6$^{th}$ cycle of spindle reassembly in *smg7-6* mutants.**

**Movie S4: Live imaging of *HTA10:RFP* chromatin marker during mitosis in a root tip of a wild type plant.**

**Movie S5: Live imaging of *HTA10:RFP* chromatin marker during mitosis in a root tip of a *cenh3-2* mutant.**

**Movie S6: Live imaging of *HTA10:RFP* chromatin marker during mitosis in a oryzalin treated root tip of a wild type plant.**

**Movie S7: Live imaging of *HTA10:RFP* chromatin marker during mitosis in a oryzalin treated root tip of a *cenh3-2* mutant.**

Figure S1

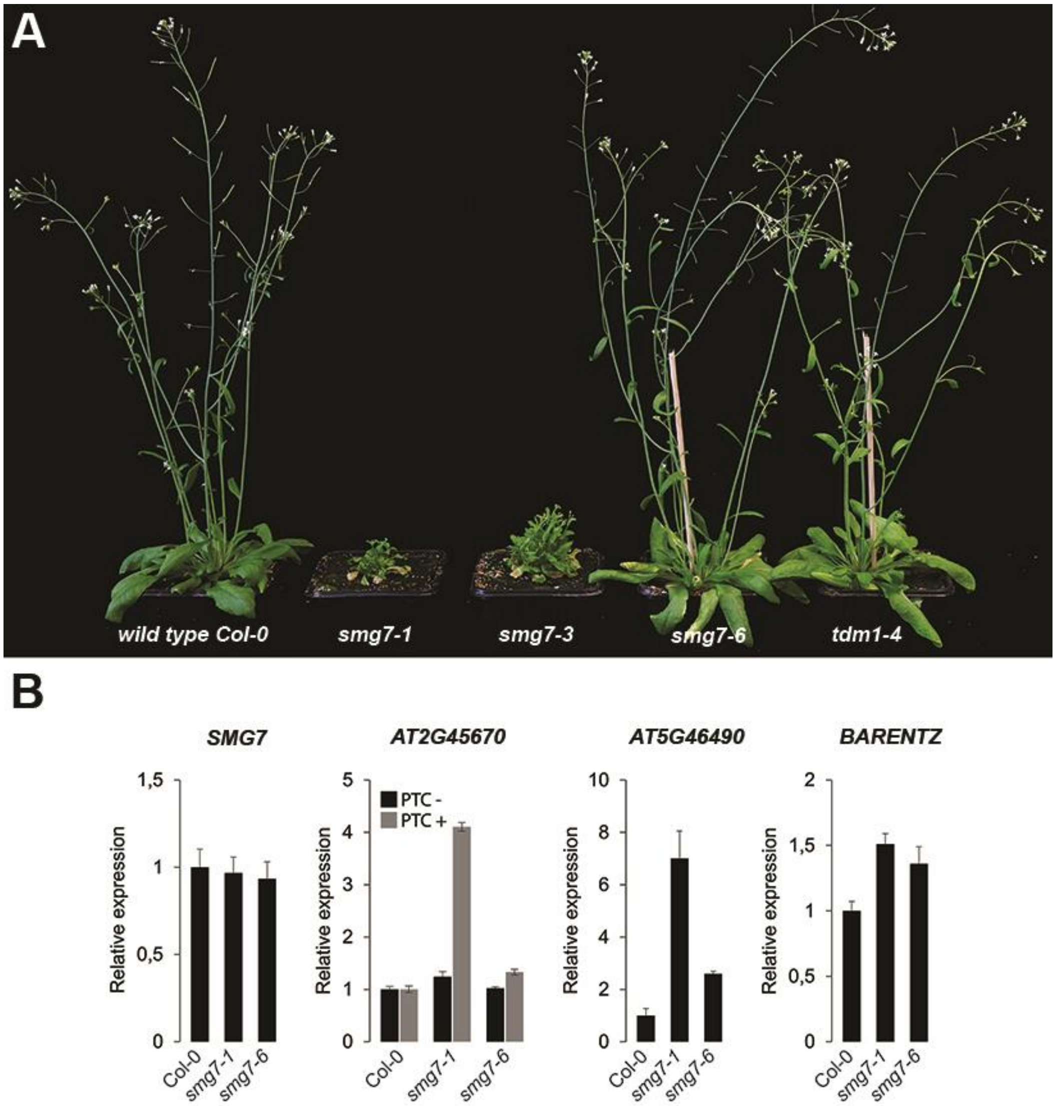


**Figure S1. Allelic series of Arabidopsis *smg7* mutants**. **(A)** Five week-old Arabidopsis mutants homozygous for the indicated alleles. **(B)** Expression of *SMG7* mRNA from the region located upstream of the T-DNA insertions in *smg7-1* and *smg7-6*. **(C)** Quantitative RT-PCR analysis of transcripts targeted by NMD in *smg7-1* and *smg7-6* mutants. Two mRNA splice variants, one containing a premature termination codon (PTC+), were quantified for the *AT2G45670* locus. Error bars indicate standard deviations from three biological replicas.

Figure S2

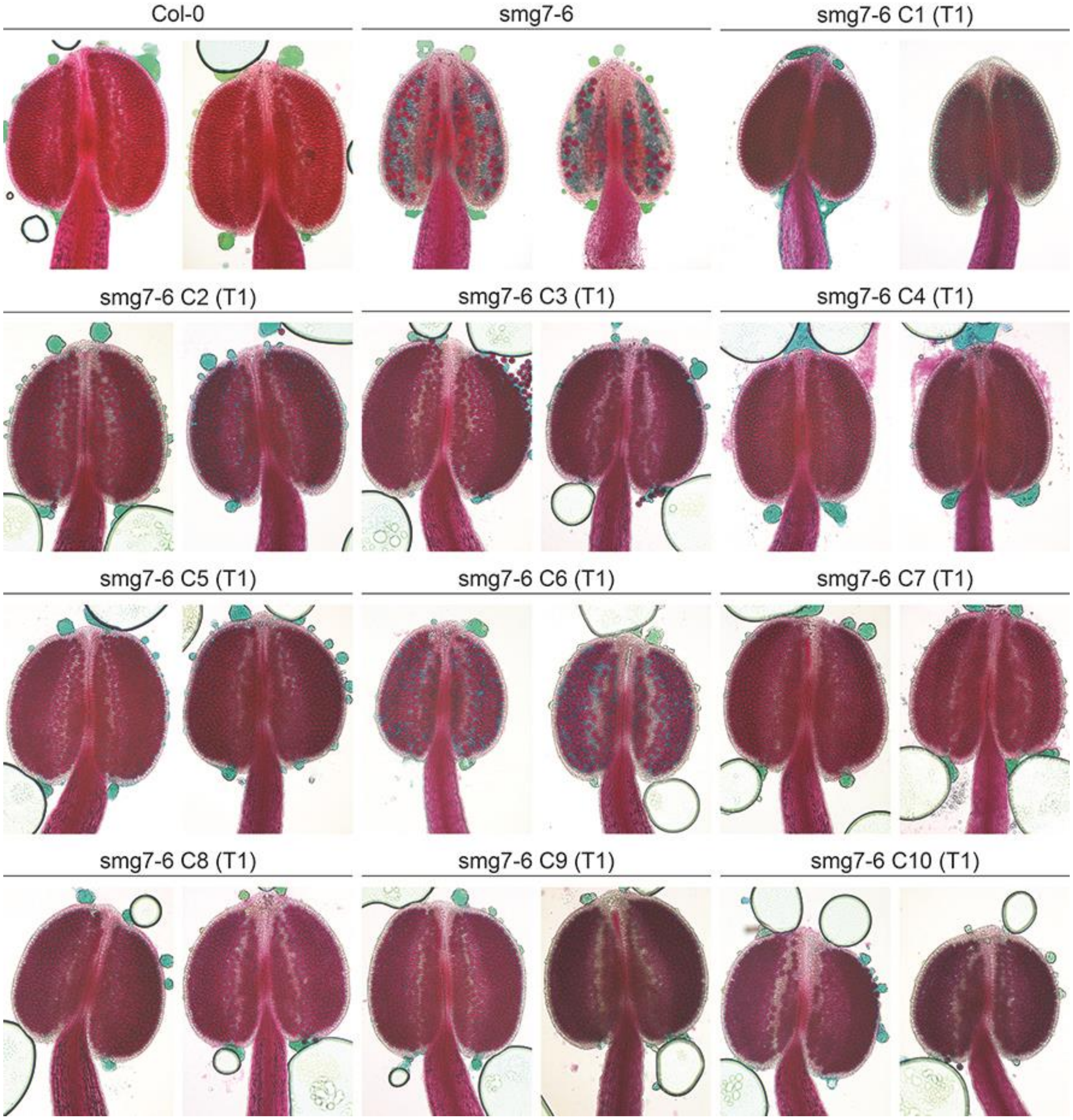


**Figure S2. Genetic complementation of *smg7-6* mutation with the endogenous *pSMG7::SMG7* gene construct.** Anthers from 10 independent T1 *smg7-6* transformants carrying the *pSMG7::SMG7* construct with viable pollen detected by Alexander staining. All T1 lines show a restoration of viable pollen.

Figure S3

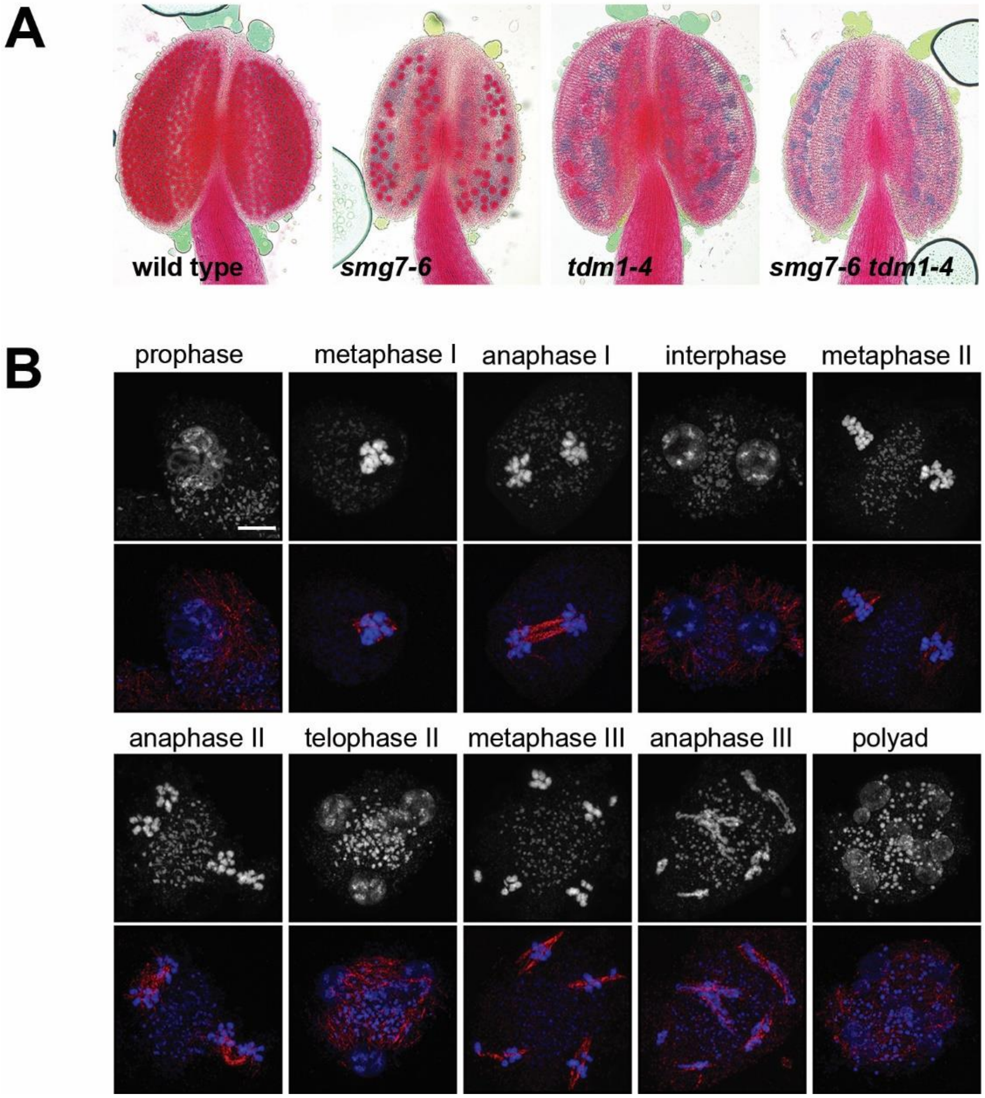


**Figure S3. Epistatic analysis of *smg7-6* and *tdm1-4* mutations. (A)** Anthers of the indicated mutants after Alexander staining. Viable pollen stain red. **(B)** Meiotic progression in PMCs of *smg7-6 tdm1-4* double mutants. Spindles are stained with anti-α-tubulin antibody (red), DNA is counterstained with DAPI. Scale bar corresponds to 5 µm.

Figure S4

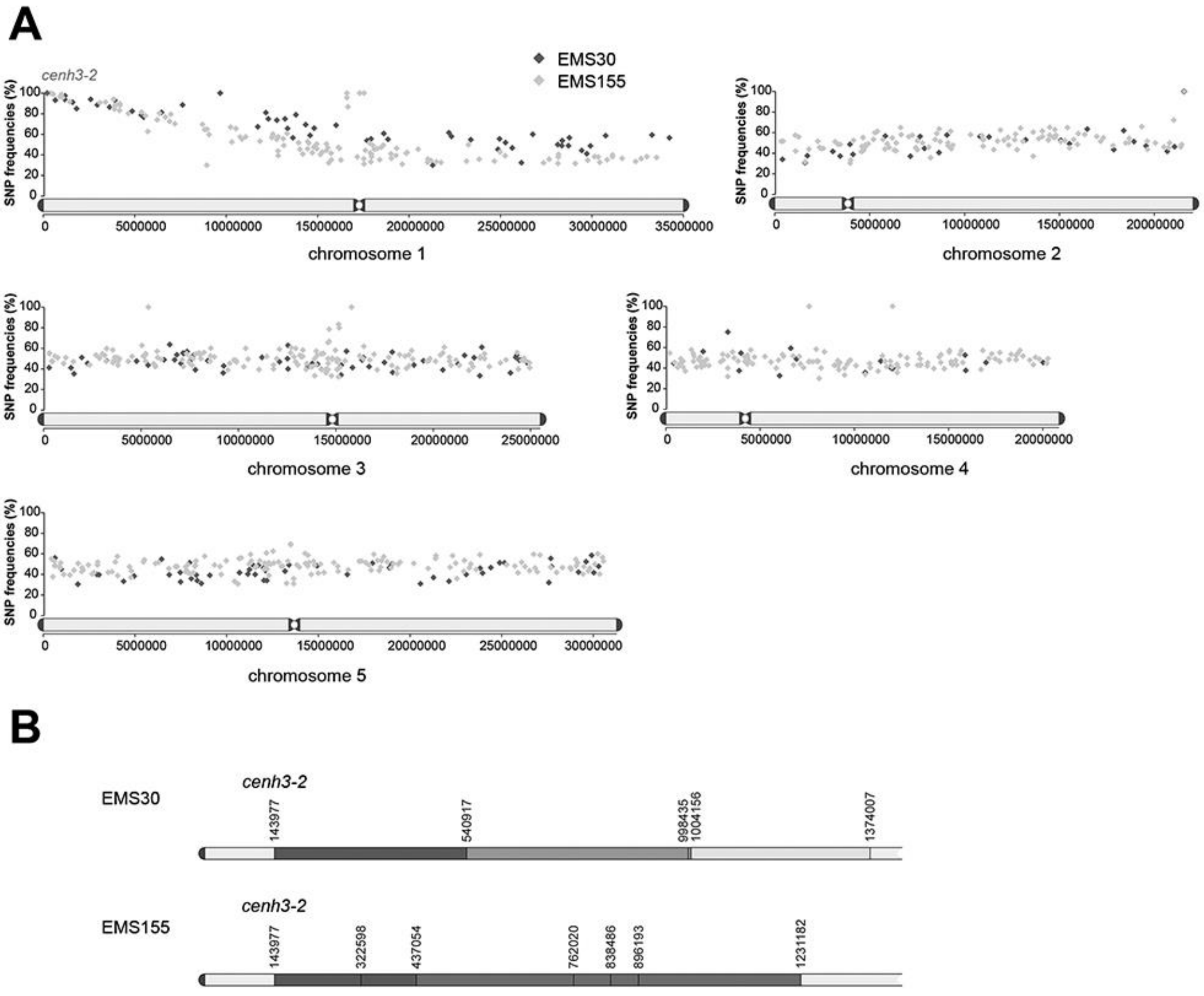


**Figure S4. Association mapping of *de novo* mutations with the *smg7-6* suppressor trait in EMS30 and EMS155 lines. (A)** Genome-wide distribution of *de novo* mutations and their frequency in fertile B2plants generated from backcrosses of EMS30 and EMS155 lines with parental *smg7-6* plants. **(B)** *De novo* mutations at the left arm of chromosome 1 in the EMS30 and EMS155 lines. Coordinates of these mutations in TAIR10 annotation are indicated.

Figure S5

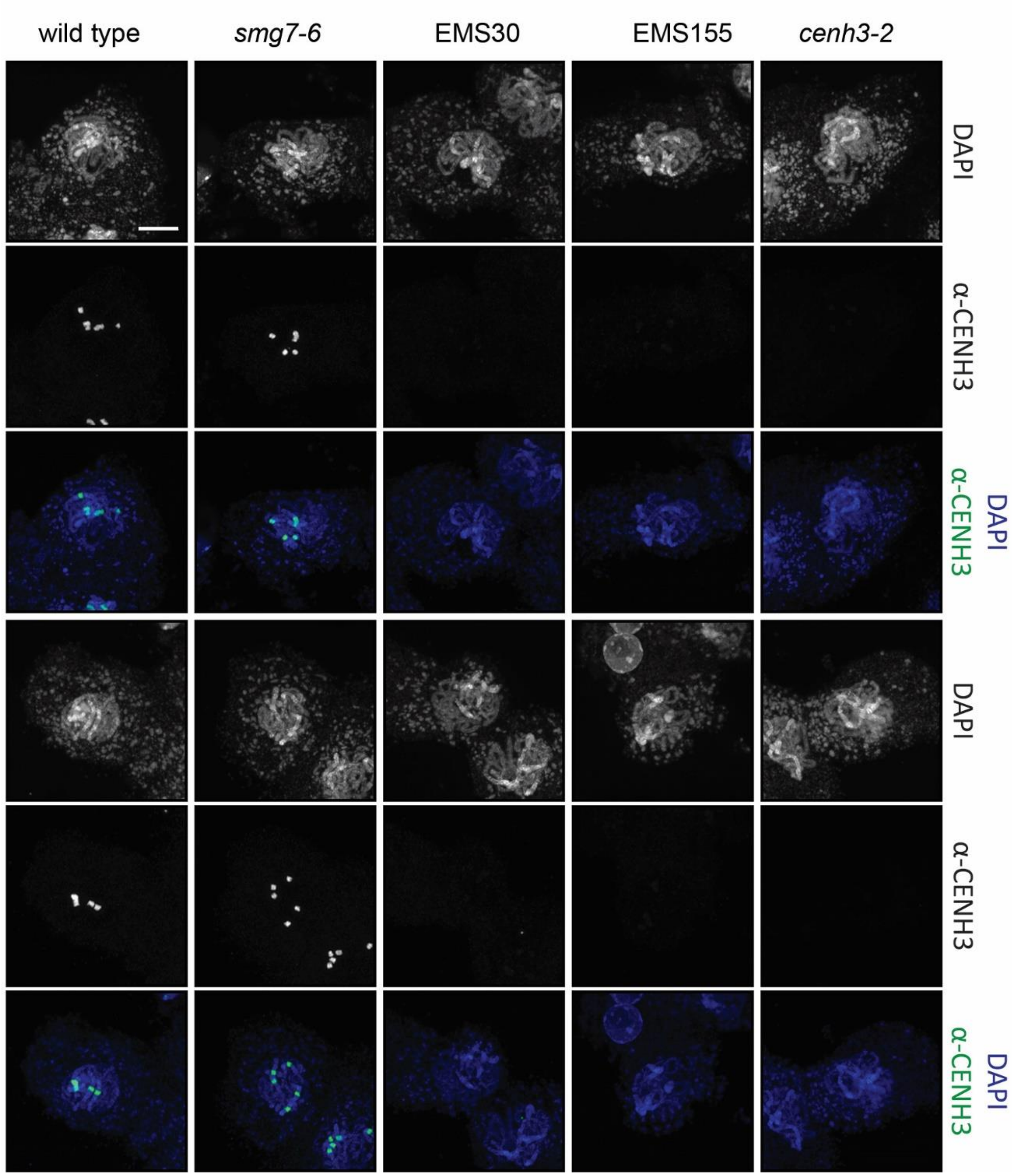


**Figure S5. Immunodetection of CENH3 in prophase I using the same imaging conditions**. CENH3 is visualized with CENH3 antibody; DNA is counterstained with DAPI (blue). Scale bar represents 5 µm.

Figure S6

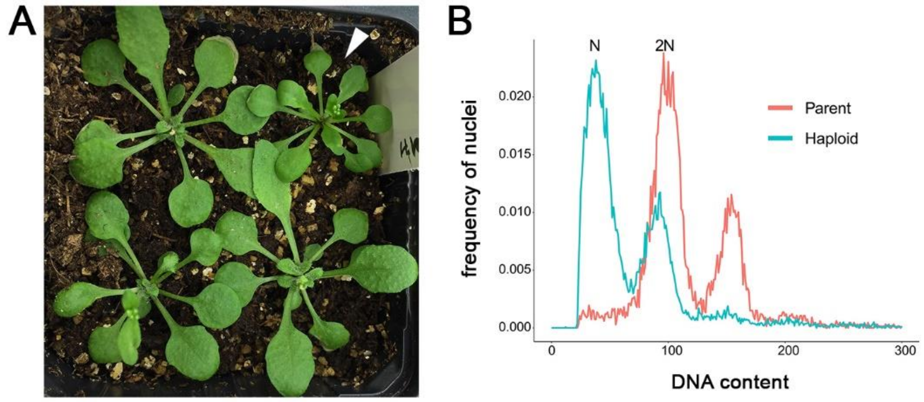


**Figure S6. Haploid plant obtained from the cross between *cenh3-2* and *gl1-1* Col-0. (A)** The haploid plant (indicated by arrowhead) was recognized based on its trichome-less phenotype. **(B)** Nuclear content of inflorescence nuclei from the haploid and a parent plant determined by flow cytometry.
